# Reconstructing disease dynamics for mechanistic insights and clinical benefit

**DOI:** 10.1101/2021.11.17.468952

**Authors:** Amit Frishberg, Neta Milman, Ayelet Alpert, Hannah Spitzer, Ben Asani, Johannes B. Schiefelbein, Siegfried G. Priglinger, Joachim L. Schultze, Fabian J. Theis, Shai S. Shen-Orr

**Affiliations:** Department of Immunology, Faculty of Medicine, Technion-Israel Institute of Technology, Haifa, Israel; Institute of Computational Biology, Helmholtz Center Munich, 85764, Neuherberg, Germany; Systems Medicine, Deutsches Zentrum für Neurodegenerative Erkrankungen (DZNE), Bonn, Germany; Department of Ophthalmology, Ludwig-Maximilians-University, Munich, Germany; Genomics and Immunoregulation, Life & Medical Sciences (LIMES) Institute, University of Bonn, Bonn, Germany; Deutsches Zentrum für Neurodegenerative Erkrankungen (DZNE). PRECISE Platform for Genomics and Epigenomics at DZNE and University of Bonn, Germany; Department of Mathematics, Technical University of Munich, 85748, Garching, Germany; Technical University of Munich, TUM School of Life Sciences Weihenstephan, 85354, Freising, Germany

## Abstract

Diseases change over time, both phenotypically and in underlying driving molecular processes. Though understanding disease progression dynamics is critical for diagnostics and treatment, capturing these dynamics is difficult, due to their complexity and the high heterogeneity between individuals. We developed TimeAx, an algorithm which builds a comparative framework for capturing disease dynamics using high-dimensional short time-series data. We demonstrate TimeAx utility by studying disease progression dynamics for multiple diseases and data types. Notably, for urothelial bladder cancer tumorigenesis, we identified a stromal pro-invasion point on the disease progression axis, characterized by massive immune cell infiltration to the tumor microenvironment and increased mortality. Moreover, the continuous TimeAx model differentiated between early and late tumors within the same tumor subtype, uncovering novel molecular transitions and potential targetable pathways. Overall, we present a powerful approach for studying disease progression dynamics, providing improved molecular interpretability and clinical benefits for patient stratification and outcome prediction.

## Introduction

Diseases are dynamic processes encompassing a multitude of changes. These range from intra-cellular molecular states, such as those occurring following cellular differentiation or activation, to changes in systemic molecular, cellular and physiological states. Identifying the underlying dynamics of diseases at high-resolution enables their quantitative comparison and is critical for designing novel preventive and therapeutic strategies to improve health. Time-series experimental designs provide an opportunity for studying disease dynamics and the variability across patients. However, while disease progression rate is patient-specific, time-series data is usually collected at fixed intervals. This reduces the efficiency of comparing progression dynamics when using time as a predictive variable and forces the clustering of data from different time points to obtain some level of shared dynamics (**Figure 1A**). Such attempts are further confounded by the fact that disease progression dynamics are often orchestrated by multiple biological processes simultaneously, requiring modeling to be performed in a high dimensional space. Dimensionality reduction methods, such as Principal Component Analysis (PCA) and Factor Analysis (FA)^1^, may be driven by time-independent variation, yielding low interpretability and making the inference of patients’ disease progression dynamics non trivial^2^. Recent advances in this area focused on the calculation of differentially expressed features over time only compared extreme states, such as disease versus controls ^3,4^, rather than describing the global dynamics of the disease. Therefore, forming a robust model for disease progression remains challenging, even when large time-series data are available.

**Figure 1:**
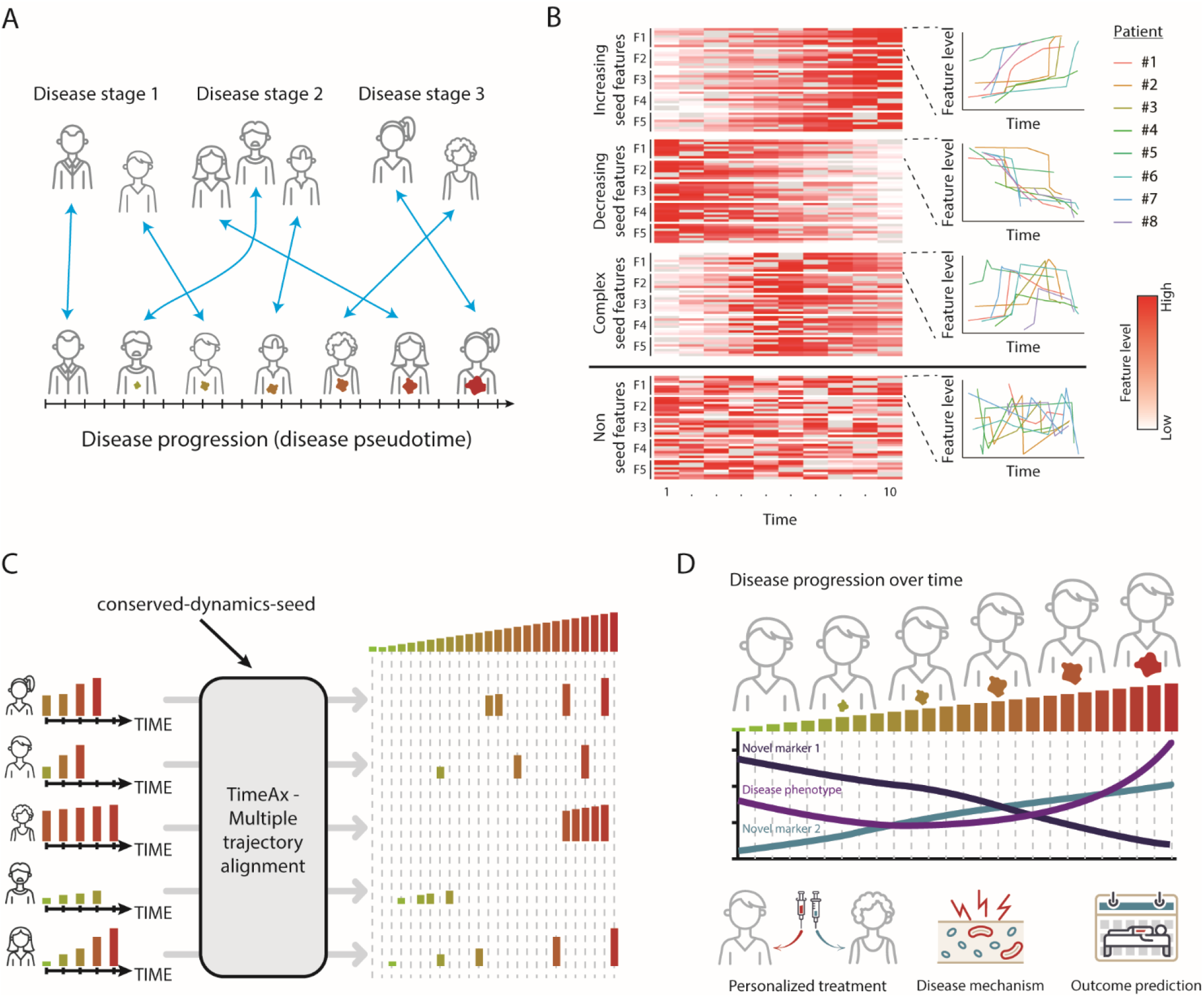
TimeAx discovers the shared disease dynamics across multiple patients. **A**. Disease progression is not captured by patient subtyping. While current patient stratification and disease state comparison requires clustering of patients into coarse subgroups (top), understanding disease dynamics as a common ground between patients’ disease trajectories enables patient stratification in a higher resolution (bottom). **B**. TimeAx seed features selection process. TimeAx selects a *“conserved-dynamics-seed”* with shared dynamics across all subjects. In the heatmap (left), each row is a patient and each column is a sample collected at a different time. Each feature spans multiple patients (multiple rows) and is also presented as a line plot (right). **C**. TimeAx captures the shared disease progression dynamics across multiple heterogeneous patients disease trajectories, based on similar concepts as multiple sequence alignment. **D**. Disease pseudotime can be utilized to discover novel disease mechanisms as well as to support new clinical frameworks for patient stratification and outcome prediction.

Here, we present TimeAx, a method that captures a representation of disease dynamics over time based on time-series data from multiple individuals. Akin to multiple sequence alignment, a historically transformative tool, which enabled biologists to build a quantitative-mechanistic understanding of DNA and protein functions ^5^, TimeAx performs multiple trajectory alignment providing major benefits for molecular interpretation and clinical diagnosis of both acute and chronic diseases. We demonstrate the utility of TimeAx for multiple time-series data types, including patients’ transcriptomes and features extracted from medical imaging. Overall, our framework allows a high resolution understanding of disease progression dynamics, discovering its underlying molecular mechanisms, and paving the way for better clinical decision making, including the design of new clinical interventions.

## Results

### TimeAx reveals shared disease dynamics across multiple patients

Patient cohorts tend to display high heterogeneity in patients’ disease courses, masking the shared disease progression dynamics and the underlying biological mechanisms shared across all patients. Often, the naive solution is clustering patients into disease clinical staging or subtypes, missing the continuous dynamics of the disease and its progression over time (**Figure 1A**). To overcome this problem, TimeAx quantitatively models a representation of the shared disease dynamics over time. TimeAx relies solely on measured features (i.e, genes, clinical markers, etc.) collected longitudinally from multiple patients (3 or more time points per patient). Patients’ time points may differ in number and in collection time. Broadly, the TimeAx process consists of three steps (**Figure S1A**, see **Methods**): First, a feature selection step, in which TimeAx uses either a user-predefined or an unsupervised, computationally-selected, set of features whose dynamics are loosely similar across patients (*“conserved-dynamics-seed”)*. While not directly associated with the hidden disease progression axis, their shared dynamics point towards their utility to serve as the backbone for the comparison between patients’ disease trajectories (**Figure 1B**). Second, TimeAx builds a model, which approximates the shared disease dynamics across all patients’ disease trajectories (**Figure 1C**). Finally, TimeAx can leverage the model to identify the disease state of a particular individual at a particular time point, referred to as ‘disease pseudotime’. In addition, TimeAx provides a robustness measure for its model, to assess the accuracy in which it captures patient dynamics over time (see **Methods**). By capturing a shared representation of disease progression dynamics across patients, TimeAx inferred disease pseudotime can simulate patients-specific disease progression states, discovering disease progression-related mechanisms and providing predictive clinical utility, otherwise missed when only following chronological time (**Figure 1D**; additional information in **Supp. Note 1**).

### Disease pseudotime captures disease progression dynamics better than chronological time

To showcase the molecular and clinical benefits of explicit disease progression modeling from longitudinal data, we contrasted disease pseudotime with chronological time as a measure of disease progression, in both simulations (see **Supp. Note 1 and Figure S1B-E**) and real-life human disease datasets, using various data types as input. We first focused on influenza infection, an acute disease which affects the immune-system over short periods of time. In this study, 17 healthy adults were challenged with influenza and then profiled longitudinally for whole blood gene expression by RNA-seq at fixed 13-15 time points within the first 108 hours following infection (**Figure 2A**, see **Methods**; total of 268 samples ^6^, denoted as ‘Longitudinal influenza cohort’). As samples were collected at fixed times, similarly across all patients, chronological time points could not be used to differentiate between symptomatic and asymptomatic patients (**Figure S2A**). TimeAx modeling of this data assembled a robust whole blood influenza infection dynamics model (**Figure S2B**; robustness score of 0.99), allowing the inference of patient-specific disease pseudotime positions. Based on the model, we observed a clear gradual increase in disease pseudotime in symptomatic but not in asymptomatic patients (**Figure 2B** and **S2C**; *p*<10^−29^), an observation which we validated using two additional cohorts, including both children and adults patients (**Figure S2D-E**; *p*<10^−47^ and 0.0003 for longitudinal adult and children cohorts respectively, see **Methods**). In addition, we identified changes in molecular processes over the disease pseudotime that would otherwise be missed using the chronological time for disease progression (**Figure 2C** and **S2F**, see **Methods**). In addition to genes which were associated with both disease pseudotime and chronological time, we identified 2274 genes which were only significantly associated with disease pseudotime, accounting for about 19% of the genes and 70% of genes with any signal. Of note, genes with positive associations with the disease pseudotime were highly enriched for multiple pathways, including the interferon and heme metabolism pathways (**Figure S2G**), in concordance with previous findings emphasizing these pathways as related to augmented immune responses ^7–10^, with implications to disease severity and patients clinical outcomes during influenza infections.

**Figure 2:**
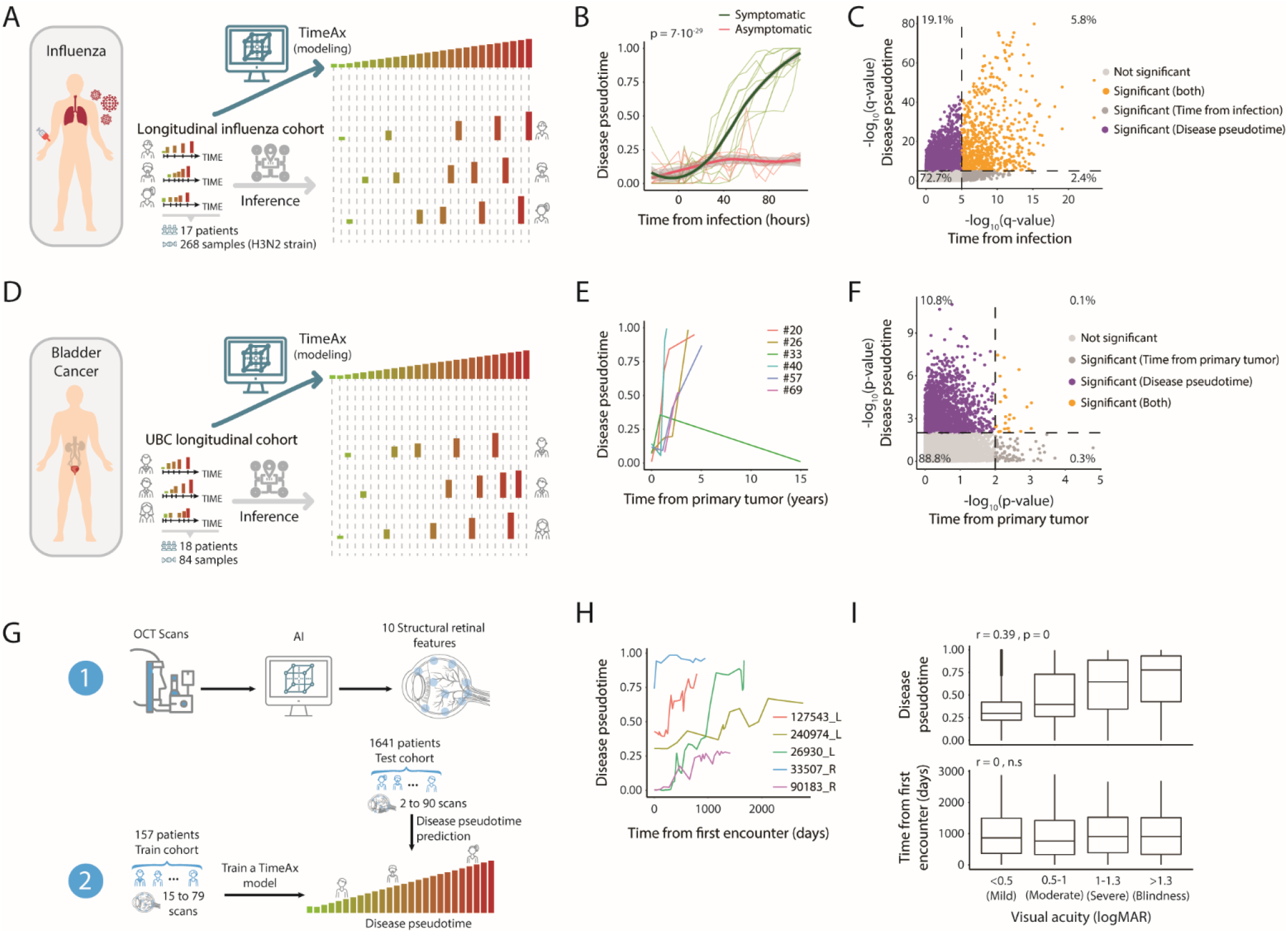
Disease pseudotime captures disease progression dynamics better than chronological time. **A**. An illustration of the influenza infection dynamics TimeAx modeling and disease pseudotime inference, based on the longitudinal influenza cohort. **B**. TimaAx disease pseudotime (*y*-axis) is different from chronological time (time from infection, *x*-axis). Shown are the differences between symptomatic (green lines) and asymptomatic (red lines) patients from the longitudinal influenza cohort. P-value was calculated using a linear mixed model. Trend lines represent the average levels in each of the groups. **C**. Gene associations (-log10 transformed, FDR-corrected, Q-values) with sampling time (*x*-axis) and disease pseudotime (*y*-axis), using a Q-value threshold of 10^−5^ (Dashed lines) (See **Methods**). Genes are colored based on their association with the two time axes. **D**. An illustration of the UBC dynamics TimeAx modeling and disease pseudotime inference, based on the UBC longitudinal cohort. **E**. TimeAx disease pseudotime (*y*-axis) is different from chronological time (time from primary tumor, *x*-axis), exemplified in six patients in the UBC longitudinal cohort. **F**. Gene associations (-log10 transformed, P-values) with sampling time (*x*-axis) and disease pseudotime (*y*-axis), using a P-value threshold of 10^−2^ (Dashed lines) (See **Methods**). Genes are colored based on their association with the two time axes, displaying significant associations almost entirely only with the disease pseudotime. **G**. An illustration of the AMD dynamics TimeAx modeling and disease pseudotime inference, based on segmented features extracted from OCT scans of the patients’ retina. **H**. TimeAx disease pseudotime (*y*-axis) is different from sampling time (time from first encounter, *x*-axis), exemplified in five patients in the AMD train cohort. **I**. The distribution of AMD test cohort patients’ disease pseudotime positions (top) and times from first encounter (bottom) (*y*-axis), across different visual severity states, determined according to the patients’ visual acuity levels (logMAR; *x*-axis). Boxes represent the 25th, 50th, and 75th percentiles; whiskers show maxima and minima.

Diseases often advance slowly and progress at different rates across patients, making it difficult to track their dynamics and understand their molecular drivers. A common approach is to cluster patient samples into disease subtypes, clinical or data-driven, incurring a loss of the continuous aspect of disease dynamics due to large differences in rates of disease progression between patients ^11,12^. To showcase the ability to capture long term processes of disease progression through explicit quantitative modeling of disease dynamics, we studied urothelial bladder cancer (UBC), a tumor with high recurrence rates after cancer removal or treatment, that mostly presents as a non-muscle invasive tumor with a small proportion of patients progressing to its muscle-invasive form, increasing the risk of developing metastases ^13^. We trained a TimeAx model using time series microarray data from 18 patients with recurring non-muscle invasive bladder cancer who ultimately progressed to advanced disease, sampled longitudinally during the incidence of tumor recurrence (**Figure 2D**, see **Methods**; ‘UBC longitudinal cohort’). In this cohort, each patient had 4-6 samples, collected up to 15 years apart from first to last recurrence^11^.

The assembled model demonstrated high robustness (**Figure S3A**; robustness score of 0.92), allowing the inference of disease pseudotime positions. These exhibited high patient variation with respect to the chronological time elapsed from their primary tumor diagnosis (**Figure 2E**). Also, disease pseudotime presented strong molecular associations, which could not be observed when modeling the data based on chronological time (**Figure 2F**; using linear regression). Specifically, we identified 2508 genes (about 11% of the genes and 96% of genes with detected signal) as significantly associated solely with disease pseudotime and not with the chronological time (**Figure 2F, upper left quarter**). These included known clinical biomarkers of UBC progression such as CCL2 and IFITM2, as well as negatively associated SGPL1, a marker linked to positive outcomes in cancer ^14–16^ (**Figure S3B**). This signal enhancement was also observed at the pathway level where a TimeAx based analysis identified stronger associations for known cancer-related processes such as the epithelial-mesenchymal transition (EMT) ^17,18^, TNFα signaling ^19^, interferon gamma ^20^ and G2/M cell cycle checkpoint ^21^ (**Figure S3C**, see **Methods**; *q*<10^−48^, *q*<10^−25^, *q*<10^−19^ and *q*<10^−9^, respectively).

Dynamic modeling by TimeAx is applicable in multiple data types, including imaging technologies commonly used for patient diagnosis and monitoring in the clinic. To further demonstrate the scope of TimeAx utility, we sought to apply TimeAx to age-related macular degeneration (AMD), a chronic disease monitored periodically using the low cost and non-invasive optical coherence tomography (OCT) ^22^. AMD is an irreversible progressive chronic disease of the retina, resulting in a decrease in visual acuity and is one of the leading causes of blindness in developed countries ^23,24^. We generated a TimeAx model of AMD progression, using segmented features, generated from OCT scans of 157 patients, each with 15 to 79 consecutive scans over the years (4953 scans overall; AMD train cohort). We then used the generated model to predict disease pseudotime positions for additional 34,836 OCT scans, collected from 1641 different patients (**Figure 2G**, see **Methods**; 2 to 90 consecutive scans per patient; denoted as ‘AMD test cohort’ ^25^). The original analysis, based on OCT scans from a subset of both cohorts, used chronological time but did not identify any changes in retinal morphology associated with disease progression ^25^. In accordance with this finding, we observed that individuals’ disease pseudotime generally increased over time, and was highly variable between patients (**Figure 2H**), suggesting that disease pseudotime accounts for patients’ variation which is only partly captured by chronological time. Indeed, we observed a strong association between the increase in disease pseudotime and the severity of the patients’ disease burden as assessed by visual acuity (**Figure 2I, top**), while no significant association was found for chronological time (**Figure 2I, bottom**). Moreover, we observed an increase, at higher disease pseudotime positions, in the usage of anti-VEGF injections, a clinical procedure for reducing the accumulation of retinal fluids which are associated with worst visual acuity (**Figure S4A**; *p*<10^−192^). This supports the notion that retinal fluids appear mostly in late stages of the disease (see **Supp. Note 2 for** full AMD progression analysis, including the identification of disease progression-related segmented features and clinical applications). Taken together, these results suggest that TimeAx dynamics modeling enables molecular interpretability and may provide increased clinical utility compared to naive longitudinal monitoring based on chronological time.

### TimeAx uncovers an advanced tumor state with unfavorable clinical outcomes

To highlight the utility of TimeAx we focused on the UBC progression model, where the long time scales of disease progression result in large heterogeneity across patients, and the high resolution molecular data enables the study of cellular and molecular mechanisms of tumorigenesis. To confirm that our TimeAx model captures tumor progression, we compared the disease pseudotime positions in the UBC longitudinal cohort with the tumor stage of the patients (see **Methods**; based on TMN staging), and saw a clear association between increased disease pseudotime and more advanced stages (**Figure 3A**, *p*<0.0005 by linear regression). We validated these results using held-out longitudinal UBC samples from the same dataset, and two additional cross-sectional UBC test cohorts -the former consisting of microarray data of 276 UBC patients ^26^ and the latter of 430 UBC RNA-seq samples from the Cancer Genome Atlas (TCGA) Program (**Figure S5A**, see **Methods**; respectively denoted as ‘longitudinal’, ‘microarray’ and ‘TCGA’ test cohorts). Tumors with higher disease pseudotime positions were also more transitional/invasive (**Figure S5B**; *p*<10^−7^), compared to more papillary tumors in lower pseudotime positions, in addition to being more common in patients with prior malignancy (defined by TCGA; **Figure S5C**; *p*<10^−4^).

**Figure 3:**
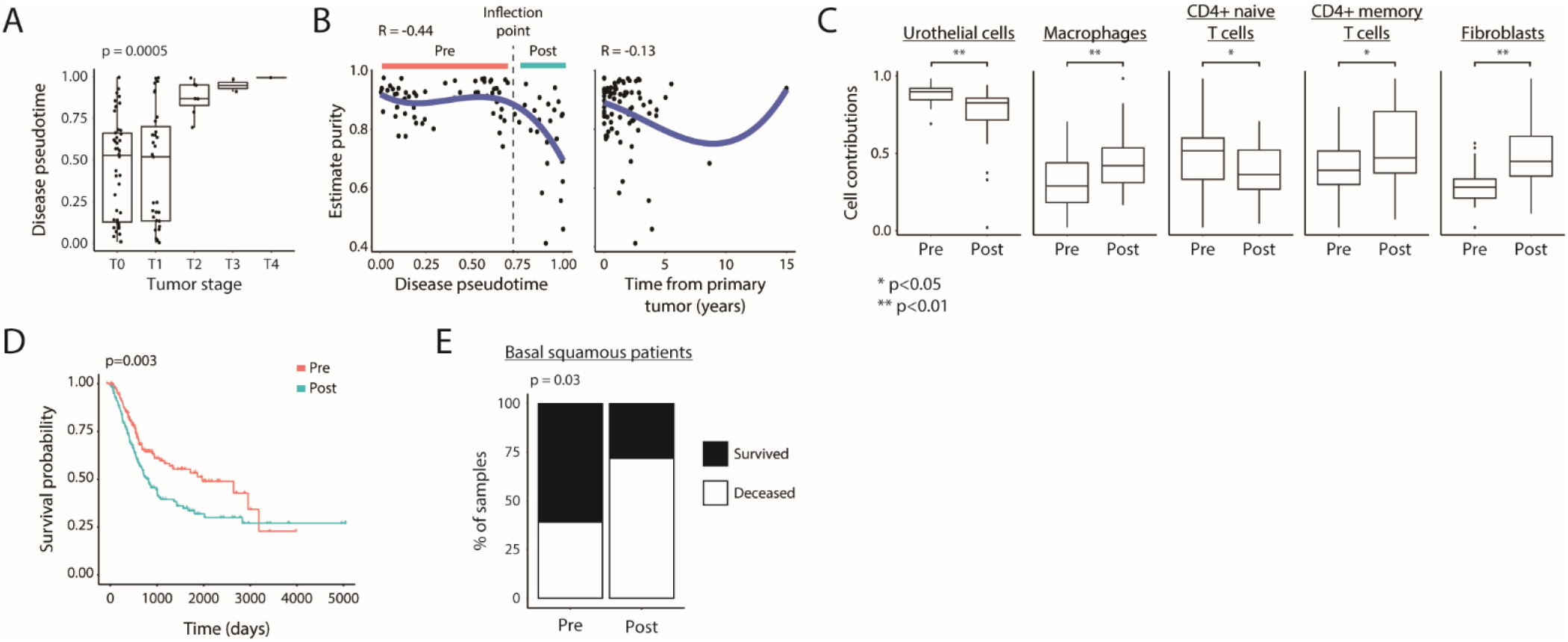
TimeAx uncovers an advanced tumor state with unfavorable clinical outcomes. **A**. Disease pseudotime clinical applicability. Shown are disease pseudotime (*y*-axis) relation with tumor stage (*x*-axis) for samples within the UBC longitudinal cohort. **B**. Tumor purity scores (*y*-axis) along the disease pseudotime (left) and the tumor recurrence times (right) (*x*-axis), displaying a sharp decrease in high disease pseudotime positions (stromal pro-invasion point; dashed line). Disease pseudotime ranges pre and post the stromal pro-invasion point are marked by colored bars. **C**. Cell type deconvolved cell contributions (*y*-axis), displaying major differences between pre- and post-stromal pro-invasion point samples. * p<0.05, ** p<0.01. **D**. Survival plot for UBC patients within the TCGA test cohort, comparing patients’ tumors with disease pseudotime positions lower and higher the stromal pro-invasion point (pre vs. post, respectively; color coded). **E**. Survival (black) versus deceased (white) rates of basal/squamous patients within the TCGA test cohort pre and post stromal pro-invasion point (*x*-axis). p-value was calculated using Fisher’s exact test. In **A** and **C**, boxes represent the 25th, 50th, and 75th percentiles; whiskers show maxima and minima.

We hypothesized that the TimeAx model, based on whole bulk-tissue transcriptomes, represents a continuous shift in complex multicellular programs in which both cell compositions and cell-states may change along the disease pseudotime, resulting in tumor progression. To explore our hypothesis, we first looked at tumor purity, a measure for the fraction of cancer cells in a tumor sample (calculated as in ^11^), along the TimeAx UBC disease progression model. We identified a position along the disease pseudotime (disease pseudotime of 0.7) where a sharp decrease in tumor purity occurred, a trend undetectable when ordering samples by the time from primary tumor occurrence (**Figure 3B**, see **Methods**; *r*=-0.44 and *r*=-0.1, respectively). A decrease in tumor purity has been previously associated with elevated immune infiltration and overall poor prognosis ^27,28^. Taken together, the TimeAx model highlights the existence of a stromal pro-invasion point, characterized by a change in the immune-stroma composition within the tumor microenvironment ^28,29^.

To understand how cellular composition and regulatory programs change along the disease pseudotime and how these relate to the decrease in tumor purity at the stromal pro-invasion point (**Figure 3B**), we deconvolved the cell composition of all samples in all four cohorts, predicting the compositions of seven major cell types: urothelial cells, muscle cells, basal tumor cells, endothelial cells and fibroblasts, as well as T cells, macrophages and additional immune cell subtypes (see **Methods**). Samples localized to disease pseudotime positions past the stromal pro-invasion point showed a significant decrease in the abundance of urothelial cells, accompanied by an increase in the abundance of activated macrophages and fibroblasts (**Figure 3C** and **Figure S5D**; *p*<0.003), a transition from naive to memory CD4+ T-cells (**Figure 3C** and **Figure S5D**; *p*<0.05) with no significant change in basal tumor cells (**Figure S5D-E**; *p*>0.1). We reasoned that differences in cell compositions along the TimeAx model may be due to either differences in disease severity as well as to technical biopsy sampling considerations. We therefore decided to leverage the clinical outcome data available in the TCGA test cohort to test whether mapping pre-versus post-stromal pro-invasion point carried clinically meaningful signal (see **Methods**). Indeed, we observed a significantly lower survival rate for patients mapping to the TimeAx disease progression axis past the stromal pro-invasion point (**Figure 3D**; *p*<0.003 by Kaplan-Meyer). The classification of patients into either pre-or post-stromal pro-invasion point, improved survival rate prediction even after accounting for other covariates with known association with survival, including age, sex and the clinical stage of the disease (**Supp. Table 1**; p<0.02, Cox Proportional-Hazards Model). Specifically, even within a specific molecular cancer subtype, such as urothelial-like and basal/squamous tumors, patients with disease pseudotime positions past the stromal pro-invasion point displayed higher mortality ratio (**Figure 3E**; 72% compared to 39%, *p*<0.03) and more rapid rates of mortality (**Figure S5F-G**). These observations suggest that the stromal pro-invasion point we observed reflects a biological milestone in UBC progression, in line with previous findings associating the infiltration of activated immune cells and cancer-associated fibroblasts into the tumor microenvironment with tumor progression and poor clinical outcome ^30,31^. Taken together, this suggests that the TimeAx model reflects tumor developmental processes and improves clinical prediction over the previously suggested tumor classification frameworks.

### Disease pseudotime captures variation undetectable by current stratification frameworks

Current clinical assessment of UBC progression, as well as the selection of interventions and therapies, relies mostly on histopathological staging. As UBC is a highly heterogeneous disease, these traditional methodologies, focusing on a relatively small set of markers, are not sufficient for optimal clinical decision making. Recently, molecular profiling analyses divided urothelial carcinomas into two major molecular types, luminal and basal, with the latter showing down-regulation of urothelial differentiation markers, being more common in muscle invasive tumors and is associated with worse clinical prognosis ^32^. The luminal type, which represents most of the non-muscle invasive tumors, can be further divided into urothelial-like and genomic-unstable subtypes, with the former harboring alterations in the FGFR3 pathway ^33–35^. While these subtyping frameworks provide a promise for patient stratification, they are not widely applied in the clinic due to their high complexity and the uncertainty of their utility for clinical prognosis over traditional frameworks ^32^. Importantly, the grouping of patients into small sets of disease subtypes results in the loss of continuity in assessing disease progression. For example, though the original analysis ^11^ of the ‘UBC longitudinal cohort’ (**Figure 2D** and **Methods**) divided patients’ UBC recurrences into distinct molecular subtypes, it also showed that clinical progression occurred in all patients whereas molecular subtyping remained predominantly stable in most patients. Moreover in some cases patients were simultaneously associated with two different subtypes ^11^. This suggests that discretized molecular subtyping does not reflect disease state, nor necessarily severity, and in some cases, dynamical changes in disease progression may be interpreted as different disease subtypes.

We hypothesized that the disease pseudotime we identified represents a generalized UBC disease dynamics axis, shared across luminal and basal molecular subtypes. Indeed, relying on previously published molecular typing (‘LundTax’ tumor molecular classification ^36^), modeling disease dynamics, while excluding patients with basal tumors, allocated all patients to nearly the same disease pseudotime positions (**Figure S6A**; *r* = 0.91), pointing out the similar molecular associations as the original model (**Figure S6B**). This suggests that these molecular subtypes reside in the same disease dynamics axis which accommodate transitions between molecular subtypes along the disease pseudotime. Supporting this, we observed higher disease pseudotime levels in more progressive molecular tumor subtypes, such as basal and mesenchymal, compared to lower values in urothelial-like subtypes (**Figure 4A**; p<10^−5^ by one-way anova). This distinction in molecular subtypes was strongly associated with patients’ allocation to either pre-versus post-stromal pro-invasion point (**Figure 4B**).

**Figure 4:**
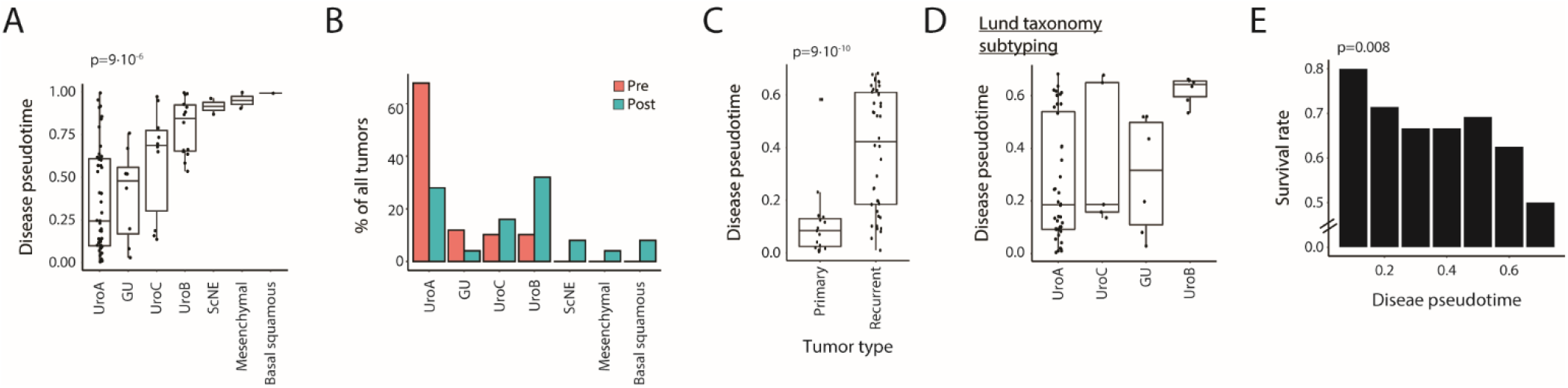
Disease pseudotime captures variation undetectable by current stratification frameworks. **A**. Disease pseudotime distribution across tumor molecular classifications in patients within the UBC longitudinal cohort based on the ‘LundTax’ molecular subtyping framework. **B**. Distribution of ‘LundTax’ tumor molecular classifications for pre (red) and post (blue) stromal pro-invasion samples within the UBC longitudinal cohort. **C**. Comparison of disease pseudotime (*y*-axis) between primary and recurrent tumors in pre-stromal pro-invasion point samples within the UBC longitudinal cohort. **D**. Disease pseudotime distribution across tumor molecular classifications in patients pre-stromal pro-invasion point within the UBC longitudinal cohort based on the ‘LundTax’ molecular subtyping framework. **E**. Survival rates of patients in the TCGA test cohort with UroA tumors (*y*-axis) across disease pseudotime bins (bin size = 0.1; *x*-axis) pre-stromal pro-invasion point. p-value was calculated based on linear regression.

We reasoned that in absence of long-term molecular follow up of patients, understanding the relation between disease progression and disease subtypes can only be garnered through further mechanistic understanding and the observation of clinically actionable predictions. As early events in UBC tumorigenesis are largely unresolved, we focused on disease pseudotime positions pre-stromal pro-invasion point. TimeAx allowed us to discern high resolution dynamics and observe differences between early (primary) and recurring tumors with a large variation in disease pseudotime between recurrent tumors (**Figure 4C**; *p*<10^−9^). Considering the present UBC molecular subtypings, we observed that while most samples were classified as low grade tumor stages (Lum-P, Lum-U by the consensus molecular subtyping ^12^ and Urothelial-like and genomically-unstable by Lund taxonomy ^36^), those patients showed large variation in disease pseudotime (**Figure 4D** and **Figure S6C**). These observations suggest that the current molecular subtyping systems of UBC tumors could benefit from the improved granularity provided by modeling the continuous dynamics of disease progression.

Of all molecular subtypes UroA exhibited the largest variation along the disease progression axis, with UroA tumors at higher disease pseudotime positions being associated with higher mortality rates (**Figure 4E**; *p*<0.01). In accordance, by dividing the UroA tumors pseudotime continuum to ‘early’ and ‘late’, respectively, using disease pseudotime cutoff of 0.25, we observed lower survival rates in late tumors (**Figure S6D**; *p*<0.1), suggesting that these tumors represent different UBC progression stages.

### Disease pseudotime uncovers novel molecular mechanisms promoting UBC progression

We next explored whether we can detect molecular patterns associated with differences between early and late progression within the UroA tumors, and whether those differences manifest different oncogenic transformation stages. We identified 2,642 differentially expressed genes between early and late UroA progression tumors, which were highly co-regulated into two main modules (*q*<0.05, **Figure 5A** and **Supp. Table 2**), one downregulated and one upregulated along the disease progression axis. In contrast, we observed no significant changes, between the two groups of UroA tumors, in the expression of established urothelial markers, including CCND1, FGFR3, FOXA1, RB1, CDKN2A, GATA3, ERBB2, PPARG and XBP1 (**Figure S6E**).

**Figure 5:**
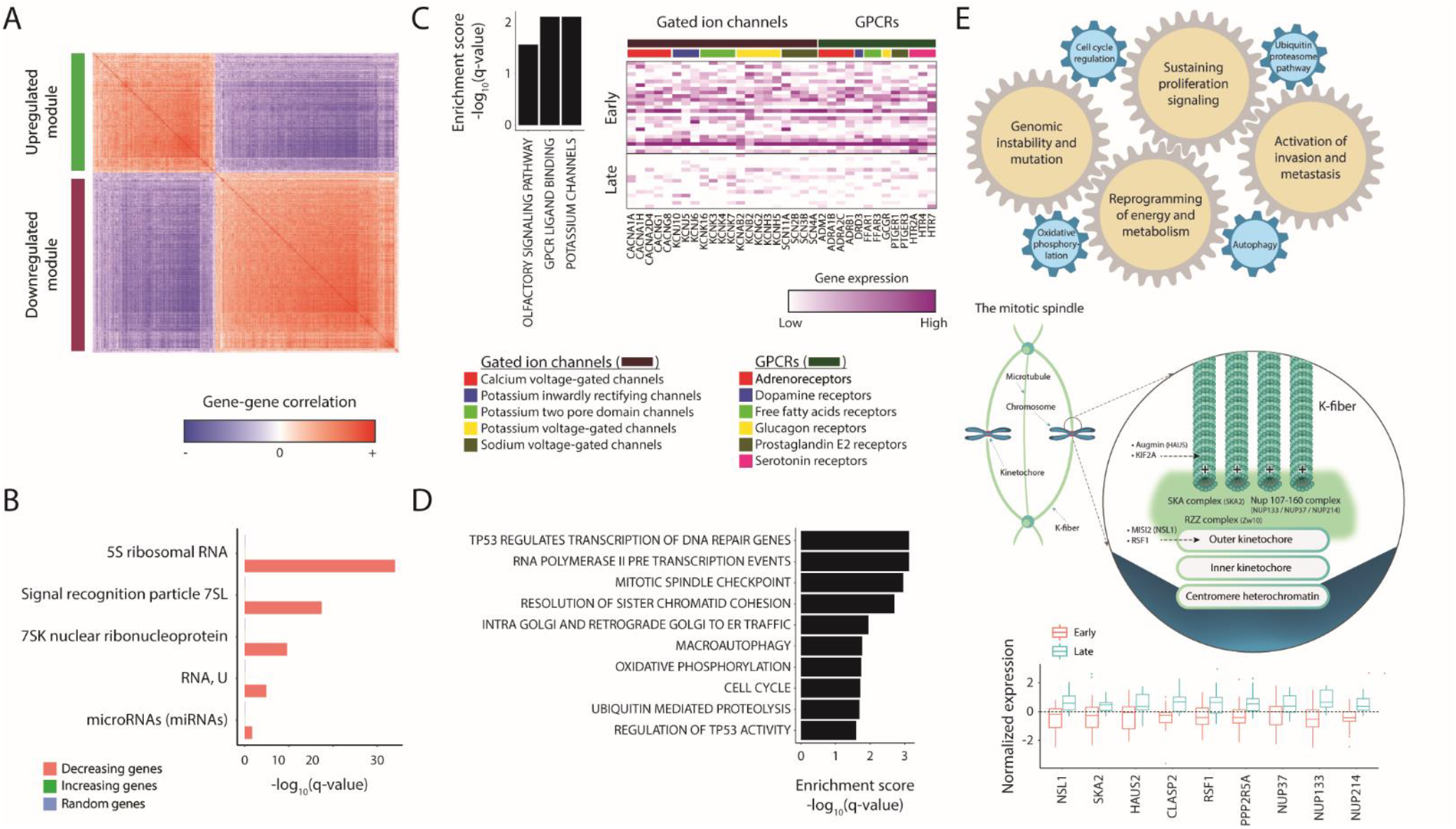
Disease pseudotime uncovers novel molecular mechanisms promoting UBC progression. **A**. Co-expression matrix for genes (rows, columns) differentially expressed (*q*<0.05) between early and late UroA tumors. Two gene sets were discovered and are highlighted in green and purple. **B**. High enrichment of different pseudogene families within the downregulated module, compared to the upregulated module set and a random gene set (color-coded). **C**. Pathway enrichment for the downregulated module, including pathway enrichment scores (left) and a heatmap of the expression levels of these pathways genes (columns), in early and late UroA tumors (rows) within the UBC longitudinal cohort. **D**. Pathway enrichment scores for the upregulated module. **E**. An illustration of the molecular model of UBC tumorigenesis, discovered by the TimeAx-based analysis, based on the increasing gene set. The illustration includes the association of known hallmarks of cancer with disease pseudotime (top) and the suggested mitotic kinetochore-spindle microtubules (MT) interaction (middle), which is also presented as boxplots of differential expression of its subunits (bottom).

The downregulated module was composed of 1587 downregulated genes within late UroA tumors (**Figure S6F**). This module was highly enriched in pseudogenes and microRNAs, (*q*<10^−4^, 0.03 respectively; **Figure 5B**, see **Supp. table 3** for functional enrichment). Interestingly, we noted that for many of these pseudogenes, their coding paralogs were transcribed by RNA polymerase III ^37^ and involved in biosynthetic processes promoting cancer cell proliferation ^38^, suggesting pseudogene actively suppress the transformation process by downregulation of their coding equivalents. In addition, we noted an enrichment for protein coding genes in this module transcribing membrane channel proteins that were downregulated along disease pseudotime. Specifically, these included ligand gated ion channels (*q*<0.05, **Figure 5C**, see **Supp. Table 4**), primarily calcium, potassium and sodium voltage-gated channels, controlling cellular ion homeostasis and downstream cell cycle and cell death processes. Interestingly, we also detected down regulation in G protein coupled receptors (GPCRs) including neurotransmitter, hormone and free fatty acids receptors (*q*<0.05, **Figure 5C**, see **Supp. Table 4**). GPCR associations with tumor progression have been unclear with evidence for them showing both tumor progressor ^39^ and suppressor ^40^ functions, likely explained both by differences in functionality between GPCRs and tissue and malignancy dependency. Our analysis showing downregulation of GPCRs in the disease progression shift towards advanced UroA pinpoints these GPCRs are likely tumor suppressors in bladder cancer.

The upregulated disease progression module contained 1055 genes (**Figure S6G**) which were highly enriched for pathways associated with malignant transformation as well as known hallmarks of cancer, including sustaining proliferative signaling, activating invasion and metastasis, genome instability and mutation and deregulation of cellular energetics (*q*<0.05; **Figure 5D**, see **Supp. Table 5**). Specifically, we detected a striking upregulation of genes in the ubiquitin proteasome (UPS) system including structural components of the 26S proteasome, components of the anaphase promoting complex and ubiquitin ligases. Post-translational polyubiquitylation of key regulatory proteins, results in their proteasomal degradation and alters regulation of cell cycle and epithelial to mesenchymal transition ^41^. In addition, late UroA tumors showed upregulation of genes functioning in cellular metabolism, including autophagy and oxidative phosphorylation, allowing the tumor to meet the increasing energetic demands and support cell proliferation during oncogenic transformation^42^.

Interestingly, we detected that the shift from early to late UroA tumor progression was associated with an abundance of genes involved in mitotic kinetochore-spindle microtubules (MT) interaction which suggested a novel mechanism for malignant transformation in UBC, undetectable without TimeAx modeling. These genes spanned six integral kinetochore constituents ^43^ which together suggest that the shift from early to late UroA during disease progression involves increasing chromosomal instability and aneuploidy. Specifically, these include MIS12 ^44^, Ska ^45,46^, RZZ ^47^ components of the outer kinetochore, components of the nuclear pore complex NUP107–160 ^48,49^ localized to the kinetochore during mitosis, the MT binding protein CLASP-2 and the chromatin remodeler RSF1 ^50^, all required for stabilizing dynamic MT to kinetochores ^51^. Similarly, late UroA tumors showed upregulation in genes coding for proteins localized to the spindle, including a subunit of the augmin complex (Haus) functioning in MT end capture ^52^ and KIF2A functioning in MT depolymerization, bi-polar spindle formation and chromosome pulling ^53^ (**Figure 5E** and **Supp. Table 2**). Taken together the signal detected via disease progression modeling versus molecular subtyping suggests that modeling of disease progression not only provides high utility for understanding mechanistic changes in the biology of disease but is likely a necessary step prior to unsupervised sub-typing of diseases.

## Discussion

We present TimeAx, a framework for studying time-dependent disease dynamics at high resolution. TimeAx modeling frees researchers from relying on chronological time in experimental designs and analyses. Instead, it makes biological time (disease pseudotime) a comparable unit that researchers can discuss quantitatively. Akin to the insight introduced by sequence alignment ^5^, and single cell cell-state trajectory inference ^54^, TimeAx opens the door to high-resolution comparative understanding of complex heterogeneous disease dynamics. Specifically, TimeAx allows the inference of a disease pseudotime position, a quantitative metric that represents the samples’ disease progression state, which can be used to build a high-resolution understanding of the temporal disease process, facilitating mechanistic discovery and patients’ outcome prediction.

We highlight TimeAx utility for both acute and chronic diseases, including the host responses after challenging healthy individuals with influenza virus, AMD disease progression and UBC tumorigenesis. In all cases, we discovered a high degree of molecular regulation, which could not be captured based on the analysis using chronological time and showcase the clinical utility of incorporating biological disease progression for patients diagnosis and disease prognosis. We demonstrate TimeAx utility for modeling diverse data types, including not only omics data, such as transcriptomics from blood or biopsy, but also incorporating data used regularly in the clinic, such as segmented features extracted computationally from OCT scans of patients over time.

Even though the underlying mechanisms of disease progression are shared across many patients, its manifestation might vary in time and magnitude within each patient trajectory, affecting the utility of chronological time as a measurement of disease progression. On the other hand, disease pseudotime presents high variation over time, capturing the hidden shared dynamics. Moreover, we note that one may use this framework to uncover patient-specific disease trajectories through repeated assembly of the trajectory, with samples from a single patient left out. Combining these trajectories with chronological time should allow the calculation of disease progression rates, providing major implications for predicting patient prognosis and patient-specific treatment.

We followed the progression dynamics of UBC recurrences, across patients over the years. Even in the expected presence of many small-accumulated environmental effects over the long lifespan of the study, we were able to devise a robust continuous disease progression trajectory, capturing the shared dynamics across all patients. The disease pseudotime, inferred by the model, supported patient stratification at a higher resolution compared to the patient stratification systems currently used in the clinic, while demonstrating improved patient outcome prediction. In addition, it captured novel cellular and molecular mechanisms of disease progression, which were hidden due to patients heterogeneity. Specifically, we observe a major drop in tumor purity at late pseudotime positions (denoted as the ‘stromal pro-invasion point’), which is associated with a remodeling of the tumor microenvironment, including a transition between naive to activated immune cells and an increase in fibroblasts. We show that patient transitions between pre-to post-stromal pro-invasion point pseudotime positions result in increased patients severity, reflecting worse patients’ outcomes, even when patients are classified among identical molecular subtypes, emphasizing the utility of the continuous disease progression model over previously suggested patient stratification frameworks.

While cell-compositional changes strongly affected the molecular profiles of the tumors, we were able to utilize the disease pseudotime to observe molecular processes which correspond to the deregulation of cellular programs associated with tumorigenesis ultimately leading to increased tumor proliferation, including a loss of cell cycle regulation and metabolic reprogramming, manifested by upregulation of genes from the ubiquitin proteasome system, autophagy and oxidative phosphorylation. Particularly interesting was the upregulation of a group of genes localized to the contact site between the mitotic kinetochore and the spindle microtubules (**Figure 5E**). The upregulation of these genes could be the mere result of increased levels of mitotic cells, due to increased proliferation rates. Another possibility, however, is the unraveling of an uncharacterized mechanism contributing to chromosomal instability (CIN) in bladder cancer. Merotelic kinetochore attachment is an erroneous process, in which a single kinetochore is attached to microtubules originating from both spindle poles during mitosis. This process is considered to be one of the major drivers of aneuploidy in mitotic cells and therefore to chromosomal instability ^55^. Correction of merotelic kinetochore attachment requires an accurately regulated rate of MT-kinetochores attachment (formation and resolution). Changes in this rate, by experimentally increasing the stability of kinetochore microtubule attachments, results in increased levels of lagging chromosomes in anaphase, indicating that slight changes in stability during mitosis are sufficient to increase chromosomal instability ^56^. UBC progression was associated with the upregulation of several genes localized to and potentially stabilizing the kinetochore-MT interaction, therefore increasing the rate of merotelic kinetochore attachment ^43^. Furthermore, the correction of erroneous kinetochore-MT interactions occurs by Aurora B kinase phosphorylating the KMN network components to reduce their affinity to MT. Our data points to the upregulation of PPP2R5A, coding for the B56 regulatory subunit of the Serine/threonine protein phosphatase 2A in UroA late tumors. This phosphatase localizes to kinetochores and stabilizes the kinetochore-MT interaction counteracting the activity of Aurora B kinase ^57^. Clearly, extensive experimental validation is required to confirm the proposed mechanism for chromosomal instability in bladder cancer. However, this sole example demonstrates the great potential of utilizing TimeAx-based disease pseudotime to provide highly predictive models for a better understanding of the molecular mechanisms leading to malignant transformation and for the discovery of potential novel drug targets.

In the present study we have demonstrated TimeAx’s utility for modeling dynamics as a one dimensional consensus trajectory, however, this does not exhaust the full potential of our framework. TimeAx can be further extended to deal with diseases displaying more complex dynamics (such as branching trajectories). Another extension of TimeAx would allow the inference of disease dynamics based on data from less than 3 time points per individual or through meta-analyses of multiple datasets. TimeAx can also be used to model other types of dynamics, such as disease recovery over time and non-disease biological processes, such as immune age ^58^. Last, while used here for gene expression and features extracted from bioimages, TimeAx can be applied to other data modalities, such as protein, microbiome and epigenetic data, as well as clinical data, including clinical markers regularly used in the clinic and multi-omic data from the same patients.

## Acknowledgements

We thank Y. Abraham for help with designing and creating figure illustrations and Martin Lukacisin and Rebecca Bendayan for their valuable feedback and discussion. This research was supported by the ISRAEL SCIENCE FOUNDATION (grant No. 1626/20), within the Israel Precision Medicine Partnership program. This work was supported in part by the German Research Foundation (DFG) to J.L.S. under Germany’s Excellence Strategy (DFG) – EXC2151 – 390873048); the HGF grant sparse2big, the EU H2020 projects SYSCID (Grant Agreement No. 733100) and ImmunoSep (Grant Agreement No. 847422). J.L.S was further supported by the BMBF-funded excellence project Diet–Body–Brain (DietBB) (grant number 01EA1809A), iTREAT (FKZ: 01ZX1902A), and by NaFoUniMedCovid19” (FKZ: 01KX2021, project acronym „COVIM”). This study was funded in part by the European Union’s Horizon 2020 Research and Innovation Program under the ERA-Net Cofund action no. 727565; the Joint Programming Initiative, A Healthy Diet for a Healthy Life (JPI-HDHL; project 529051018) and under the ERA-CVD non-cofunded action JTC2017 (Mechanisms of early atherosclerosis and/or plaque instability in Coronary Artery Disease)) awarded to J.L.S. The results here are in part based upon data generated by the TCGA Research Network: https://www.cancer.gov/tcga.

## Author contributions

SSO conceived the idea, AF and SSO developed the TimeAx method, NM led the biological interpretation, AA and FJT contributed to method development, AF performed the analysis. HS, BA, JBS, SGP and FJT contributed to the image analysis. JLS helped with biological interpretation. AF, NM and SSO wrote the manuscript and all authors reviewed and revised it.

## Code availability

TimeAx is publically available as an R package at Github: https://github.com/amitfrish/TimeAx.

## Online Methods

### The TimeAx algorithm

TimeAx aims to model the entirety of a disease’s dynamics and construct a quantitative framework through which one can better compare individuals’ sample states as part of a dynamic process shared by all individuals under examination. TimeAx takes as input a compendium of annotated measurement profiles with each profile describing a sample and a time point assayed when the sample was undergoing a biological condition whose trajectory we are interested in delineating. Profiles are a quantitative snapshot of the abundance of different measured data types, for example ‘omic’ features (e.g. genes or proteins).

The TimeAx algorithm can be divided into three steps: First, a feature selection step, in which it selects a set of seed features with shared dynamics across different individuals. Second, an alignment step, in which a consensus trajectory is being composed. Finally, a prediction step, in which different profiles are fitted to the consensus and their specific disease pseudotime position is projected.

### Step 1 (conserved-dynamics-seed selection)

Identifying the consensus of a disease requires common ground. We thus start off by choosing a set of *“conserved-dynamics-seed” features (seed -features)*. These features can be predefined by the user or can be computationally selected by focusing on features *whose* dynamics are similar across individuals. Features are scaled into [0,1] range by subtracting their minimal value and dividing by the maximal value minus the minimal value. To avoid overfitting, TimeAx focuses on those features with high variation across the profiles. Therefore, it requires seed features to have many unique values (different values in most of the measured profiles) and a high standard deviation across profiles, both of which must be above user determined thresholds. Next, for each feature, TimeAx applies an iterative bootstrap-like process, where in each iteration it samples an equal number of time points from each pair of patients and calculates their spearman correlation for the feature’s expression across these time points. Following this, TimeAx averages correlation values across all pairs of patients and iterations and chooses *L* (user-defined; default of 50) features with the highest values as being part of the “*conserved-dynamics-seed”*. Alternatively, the “*conserved-dynamics-seed”* can be pre-defined as an input by the user, disabling the computational selection of seed features.

### Step 2 (multiple trajectory alignment)

To assemble a consensus trajectory describing the complete disease dynamics, we rely on principles stemming from the mature field of sequence alignment. Specifically, in progressive multiple sequence alignment (MSA) methods, sequences are added into the MSA in an orderly fashion based on a guide tree, structured according to the pairwise alignment distance of all sequences ^5^. Similarly, TimeAx performs a multiple trajectory alignment (MTA) on time-series datasets of individuals each of which we consider as an individual partial trajectory. The alignment process relies solely on the above described “*conserved-dynamics-seed”* (Step 1).

In TimeAx, the leaves of the guide tree represent the original individual subject partial trajectories, whereas the inner nodes of the tree represent pseudo-trajectories created based on different combinations of the leaves’ trajectories. During the multiple alignment step, TimeAx follows the structure of the tree and merges each two child nodes into a new parent node. Specifically, given trajectories 1 and 2, TimeAx leverages dynamic time warping ^54,59^ to detect the best fit between sample profiles in trajectory 1 and in trajectory 2, keeping the order of both trajectories. Based on this pairwise alignment, TimeAx yields a new, intermediate trajectory with a confidence score, calculated as the average of correlation coefficients between the aligned profiles that were used in its creation. Next, TimeAx calculates new intermediate profiles as the average between the aligned profiles, weighted by their confidence scores. The confidence scores of the initial trajectories (corresponding to different individuals) are the average of the confidence scores obtained by aligning them with all the other trajectories. At the end of the process, the final node, located at the root of the guide tree is the consensus trajectory.

#### Pseudocode

Let *trajectories* be a set of *n* trajectories, each represents a time series data from an individual.

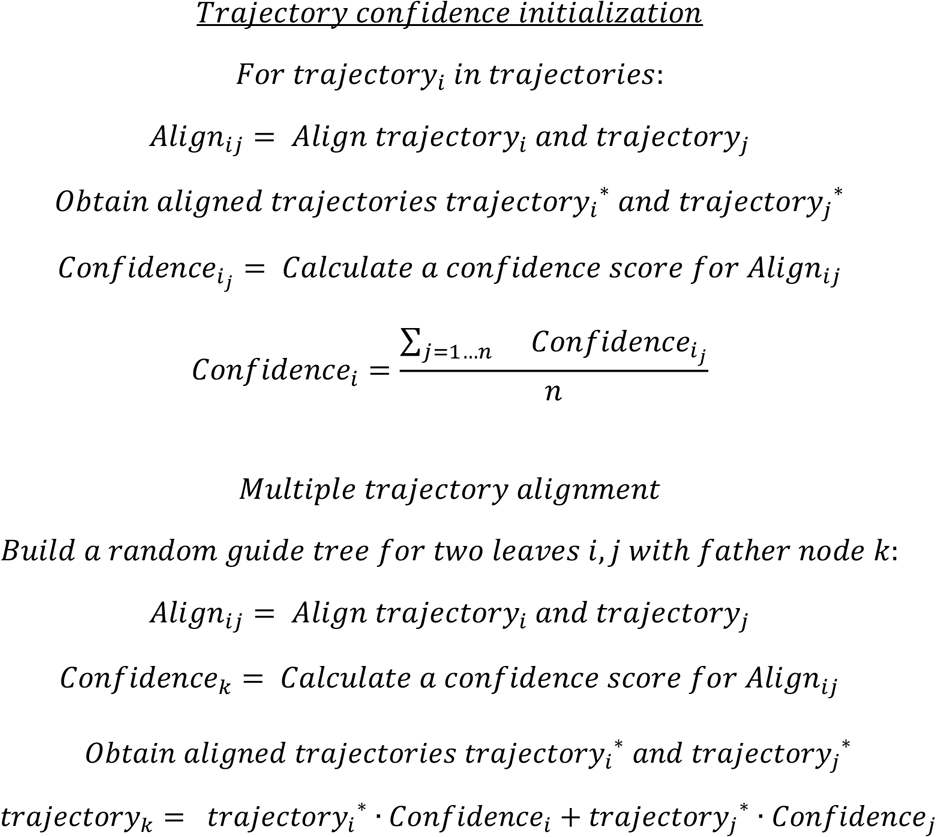

Repeat the above process for each two objects with a shared father node, until reaching the root of the guide tree.

Define the consensus trajectory as the *trajectory*_*root*_

To obtain a continuous measure of progression across the consensus trajectory, TimeAx then calculates numeric disease pseudotime positions for each of the consensus trajectory profiles. To do so, we apply a principal component analysis (PCA) to the profiles (using only the seed features), and, beginning from PC1, the set of PCs that sum up to 90% of the variance are selected. The sum of distances across all selected PCs are then calculated and transformed into a [0,1] scale, by removing their minimal value and dividing by the maximal value minus the minimal value.

One of the obstacles of MSA is the selection of a good guide tree. As the structure of the guide tree determines the alignment ordering, it has a strong effect on the resulting consensus trajectory. To overcome this, TimeAx takes an ensemble approach, creating N random guide trees (along this manuscript we use N=100 by default), representing a set of weak learners ^60^. Therefore, while each guide tree is non-optimal, the combination of the whole set provides high accuracy and robustness levels. Based on this selection of guide trees, the final model obtained by TimeAx, contains multiple consensus trajectories which can be used to predict the disease pseudotime of the original samples across individuals, as well as allow the disease pseudotime prediction for new samples (Step 3).

### Step 3 (disease pseudotime prediction)

TimeAx allows the prediction of disease pseudotime for new samples, based on the set of consensus trajectories, generated during the alignment process (Step 2). To overcome technical differences between the consensus profiles, devised based on the train longitudinal data, and the new samples, TimeAx allows the prediction to be based on the ratios between features rather than the features themselves. Regardless, TimeAx first normalizes the values of each feature within the consensus profiles and uses the normalizing factors (mean and standard deviation) to transform the new samples as well. Next, TimeAx calculates the spearman correlation between each new sample normalized profile and the normalized profiles of all states within the consensus trajectory and chooses the highest correlated state as its predicted local disease pseudotime position. Finally, as an ensemble method, TimeAx calculates the disease pseudotime position by averaging the predictions across all guide trees. TimeAx also provides an uncertainty score for each sample profile, by calculating the standard deviation across the predicted disease pseudotime positions. By definition, low value of uncertainty implies a high concordance between disease pseudotime positions across all consensus trajectories.

#### Trajectory robustness score

We devised a robustness score to assess the success of capturing a meaningful dynamics of the disease modeled by TimeAx, under the assumption that on average the disease dynamics in patients progresses towards some end goal (e.g. disease resolution, progressive disease, or some new homeostatic state). The robustness score reflects the level of agreement between TimeAx derived disease pseudotime and a disease pseudotime, assembled with a constraint that the disease pseudotime for consecutive samples within the same individual cannot be lower than previous disease pseudotime (see below). Assuming the TimeAx model is robust, consecutive profiles should on average advance along the consensus trajectory, and therefore should provide similar predictions to the constrained model, even though sample ordering was not provided as input. On the other hand, if predictions largely differ, it implies that TimeAx failed modeling the change in condition over time (or that the disease does not obey the underlying disease progression assumption TimeAx relies on). In that case, the model robustness will be low.

To derive the robustness disease pseudotime from the consensus trajectory, for a new individual with L profiles, TimeAx first calculates distance matrix D, as one minus the spearman correlation between its profiles and the M profiles of a consensus. The distances are then smoothed to increase prediction robustness (smoothing as in CellAlign interpolation ^54^). Next TimeAx constructs a weighted directed graph with L layers and M nodes in each layer. Each node in layer i is connected to nodes in layer i+1 that represent profiles located in the same level or higher in the consensus trajectory. The weight of an edge that is directed to node j in layer i is D_ij_ [*i* ∈ {1 … *L*} &*j* ∈ {1 … *M*}]. In addition, a “start” node is directed to all nodes in the first layer and all nodes in layer L are directed to an “end” node (**Figure S1F**). Finally, Dijkstra’s algorithm is used to calculate the shortest path between “start” and “end”, pointing out the best set of states in the consensus fitting to the profiles of the new individual. To obtain the final robustness disease pseudotime, the results across all different computed consensus trajectories are then averaged. The robustness score is calculated as the Pearson correlation coefficient between disease pseudotime and robustness disease pseudotime.

#### Synthetic data analysis

To create synthetic datasets that simulate the dynamics of a biological condition progressing through time, we first defined a biological trajectory and projected multiple partial trajectories, each containing multiple samples, across it (using the ‘splatSimulate’ function in the ‘splatter’ R package). Each sample profile was modeled as the expression value of 1,100 genes, 100 of which contained trajectory relevant signals and the rest not. Specifically, the 100 signals containing genes were directly associated with the biological trajectory across all simulated samples. The expression value of the 1000 non-signal genes was set as randomized values obtained by shuffling the values of the signal containing genes, across the trajectory. We then added technical noise to the data, ranging from almost-zero noise to entirely non-informative data.

For each simulation setup we then generated 100 individuals, each containing *m* consecutive samples (ranging from 3 to 20) across the trajectory. For each individual, we chose a center point within the biological trajectory and picked *m* specific points, corresponding to samples, distributed normally around the center point. We denoted the exact disease pseudotime that corresponds to the selected samples as *“Simulated disease pseudotime”*, whereas the *“Simulated time points”*, are the samples’ order within each individual (1…m). Applying TimeAx in this case implies that given an input matrix of the molecular features profiled per person, annotated with the ‘original time points’, TimeAx predicts the corresponding disease pseudotime positions, denoted as *“TimeAx disease pseudotime”*.

To test the utility of the *TimeAx* model, we devised two different types of tests: (1) To study TimeAx’s ability to accurately predict the correct disease pseudotime position for each individual we calculated the earth mover’s distance (EMD) between the *Simulated disease pseudotime* and the TimeAx *disease pseudotime* for each individual and compared it to the same distance calculated based on the *Simulated time points* instead of the TimeAx *disease pseudotime*. The final distance scores, in both cases, are calculated by averaging the values across all individuals.

(2) To contrast the ability to discover new biological signal in a naïve time series modeling approach versus TimeAx, we calculated false discovery rate (FDR) corrected p-values (denoted as Q-values) for the linear regression models using the *Simulated time points* versus the TimeAx *disease pseudotime* as predictors of the genes’ expression values across all samples. We then selected genes with Q-values of less than 0.01 as associated with progression. As only the first 100 genes of the synthetic data were truly associated with the *simulated disease pseudotime*, we assessed the gene detection efficiency based on the F1 score for each dataset.

#### Public datasets

For influenza dynamic modeling, we trained the TimeAx model using data from an influenza challenge study (H3N2/Wisconsin strain) in which 17 healthy volunteers, between 18 to 45 years of age, were infected and then profiled longitudinally over 13-15 timepoints (a subset of 0, 5, 12, 21, 29, 36, 45, 53, 60, 69, 77, 84, 93, 101 and 108 hours after infection) for whole blood gene expression by RNA-seq (total of 268 samples). All patients were treated by oseltamivir orally on a daily basis along the study ^6^ (Longitudinal influenza cohort; accession number GSE30550). To ensure that we capture disease progression dynamics, we trained the model using only the symptomatic patients (9 patients) but inferred disease pseudotime for all patients. We then validated our results using an additional data from a challenge study of an influenza infection (H1N1 strain) in 24 human adults, aged 20-35 years old, with similar study settings as in the longitudinal influenza cohort ^61^ (total of 382 samples; Longitudinal test cohort; accession number GSE52428) and a blood microarray data from healthy and H1N1 infected children (n=19, 33 respectively), all below the age of 17 ^62^ (Children test cohort; accession number GSE42026).

For the UBC disease progression model, we trained TimeAx using a time series microarray data of microdissected tumors collected from 18 recurring non-muscle invasive bladder cancer (NMIBC) patients (3 males and 15 females), at different events of tumor recurrence (4 to 6 samples per patient, collected up to 15 years apart from first to last recurrence). At least one full induction course of intravesical Bacillus Calmette-Guerin (BCG), an immunotherapy for early-stage bladder cancer, was given for some of the patients ^11^ (UBC longitudinal cohort; accession number GSE128959). To accurately capture the variation between tumor samples, we used a conserved-dynamics-seed containing 100 genes. The results for the UBC longitudinal cohort were validated in additional 28 patients (22 males and 6 females; total of 52 samples), which were excluded from the UBC longitudinal cohort due to having less than 4 samples per patient (denoted as ‘longitudinal test cohort’) and two additional test cohorts: a microarray gene expression data from the bladders of 276 UBC patients, who underwent radical cystectomy, not receiving any treatment before bladder extraction ^26^ (microarray test cohort; accession number GSE83586) and RNA-seq data from 430 UBC patients (116 females and 314 males), aged 34-90 with a median of 69, from the Cancer Genome Atlas (TCGA) Program (TCGA test cohort; downloaded from the TCGA data portal).

For AMD dynamic modeling, we trained TimeAx using conserved-dynamics-seed of 10 previously segmented features (without conserved-dynamics-seed selection), including retinal atrophy, fibrosis, retinal thickness, epiretinal membrane, neurosensory retina, subretinal hyperreflective material, retinal pigment epithelium (RPE), fibrovascular PED, drusen and choroid, derived from OCT scans of 157 patients, each with 15 to 79 consecutive scans over the years (4953 scans overall; AMD train cohort) ^25^. Using this model, we predicted disease pseudotime for additional 34836 OCT scans collected from 1641 patients (2 to 90 consecutive scans per patient; AMD test cohort), using the same segmented features ^25^. In this dataset, patients were either not treated or treated by eye injections of anti-VEGF agents. Additional features, including posterior hyaloid membrane and intraretinal and subretinal fluids, were excluded from the model due large technical variation and the direct effect of treatment.

#### Cell type deconvolution and single cell analysis

We inferred cell type compositional abundance by applying Cibersort ^63^ on UBC bulk gene expression profiles, using two different sets of signature profiles. The first set contained cell type profiles obtained directly from the UBC tumors and their microenvironments. Specifically, based on single cell RNAseq data from a muscle-invasive urothelial bladder cancer patient ^64^, we calculated the mean expression profiles of seven major different cell types: Urothelial cells (88 cells), T cells (369 cells), Muscle cells (186 cells), Basal tumor cells (436 cells), Endothelial cells (362 cells), Macrophages (283 cells) and Fibroblasts (273 cells). This set allowed us to focus on changes in the abundance of non-immune as well as the main immune cell subsets. To study higher resolution immune cell subsets, we used the LM22 reference matrix ^63^, which contains gene expression profiles of different subtypes of macrophages, T, B, NK and dendritic cells.

In addition, we focused on basal tumor cells within the single cell data and imputed the main two axes of variation, based PCA analysis using the top 1000 highly variable genes (HVGs). We linked these axes to tumor progression by presenting the association between the mean expression of HVGs and that of epithelial-mesenchymal transition (EMT) genes, observing similar spread across the two axes.

#### Gene associations with disease pseudotime

For each cohort in this study, gene associations with disease pseudotime were calculated as ‘Pearson’ correlations between gene expression levels and the disease pseudotime positions across all the cohort’s samples. To compare the number of disease pseudotime vs. sampling time associated genes, we calculated the FDR-corrected p-value (Q-value) based on a linear regression between gene expression and either of the time axes using a single Q-value cutoff (10^−5^ and 0.01 for **Figure 2C** and **2D**, respectively). In **Figure S2E**, we present a generalized comparison across multiple Q-value cutoffs (q<10^−5^, 10^−4^, 10^−3^, 10^−2^, 10^−1^).

#### Pathway enrichment

To explore the relations of biological functions with the progression of the disease, we tested candidate pathways from the following sources: GO, MSigDB Hallmark gene sets, Reactom, and KEGG. We correlated the expression levels of each gene with the inferred disease pseudotime across samples (Pearson’s r score). Next, for each pathway, we calculated a significance score for the difference in distributions between correlation coefficients of pathway member genes versus background genes that are not members of the pathway, using a Kolmogorov–Smirnov test (KS test). Finally, to obtain enrichment scores and allow their comparison across pathways, we applied a –log10 transformation.

