## Supplementary materials for "Reconstructing disease dynamics for mechanistic insights and clinical benefit"

### Supplementary Note 1

To test TimeAx's ability to capture the dynamics of a disease, we first performed a simulation study. We simulated bulk gene expression measurements from different individuals sampled at multiple time points and undergoing a common biological process with different dynamics (see **Methods**). In this scenario the disease pseudotime position on the consensus trajectory of each simulated sample is known. Next, we tested the correlation between simulated disease pseudotime (i.e. gold standard) and the simulated time points and TimeAx predicted disease pseudotime. TimeAx predicted disease pseudotime highly overlapped with the simulated disease pseudotime within each individual, across different levels of added noise, while the simulated time points failed capturing the heterogeneity between samples (**Figure S1B,** see **Methods**). One important result of TimeAx alignment is an improved ability to discover genes whose expression is closely related to disease progression dynamics (See **Methods**; **Figure S1C**). The accuracy of disease pseudotime inference increased when more samples were available per individual (**Figure S1D**). Finally, the number of consensus trajectories had only a marginal effect on TimeAx's efficiency, as five consensus trajectories were often sufficient (**Figure S1E**).

**Supplementary Note 2 - TimeAx deciphers AMD disease dynamics from longitudinal monitoring of clinical imaging data**

We studied how structural changes within the retina are associated with the progression of AMD. We discovered a strong positive correlation of disease pseudotime with retinal fibrosis and atrophy, as opposed to a negative correlation with retinal pigment epithelium (RPE) density, in agreement with previous findings of these structural features as hallmarks of AMD progression [1–4] (r=0.47, 0.7 and -0.68, respectively; **Figure S4B**). The progression along these structural features followed a nonlinear fashion, with a relatively low degree of change in low disease pseudotime positions, compared to a major shift at high positions, suggesting the initiation of a more aggressive state of the disease (at disease pseudotime position 0.75; **Figure S4B**). In addition, we found a peak in drusen at low disease pseudotime positions (at disease pseudotime position 0.25), corroborating drusen as an early marker of AMD [5] (**Figure S4B**). Finally, we observed a gradual decrease in the density of neurosensory nerves in the retina density over the disease pseudotime, pointing to a novel association with disease progression, presenting distinct densities throughout all stages of the disease (**Figure S4B**).

We wanted to showcase the utility of disease pseudotime for clinical diagnosis of AMD patients' disease states. Since the TimeAx model integrates the contributions of different structural features of the retina, we hypothesized that it will capture an improved representation of the clinical state of the patients. Indeed, different visual acuity levels were characterized by distinct ranges of disease pseudotime positions, while only retinal atrophy displayed distinct levels at higher visual acuity, showing no difference at earlier visual acuity levels (**Figure S4C**). This highlights the benefits of monitoring early disease progression stages, where disease progression can still be hindered, using disease pseudotime. In addition, by dividing patients into severity groups based on distinct levels of features, we found the less false positive classifications, based on disease pseudotime, supporting its utility for patient diagnosis and monitoring (**Figure S4D**).

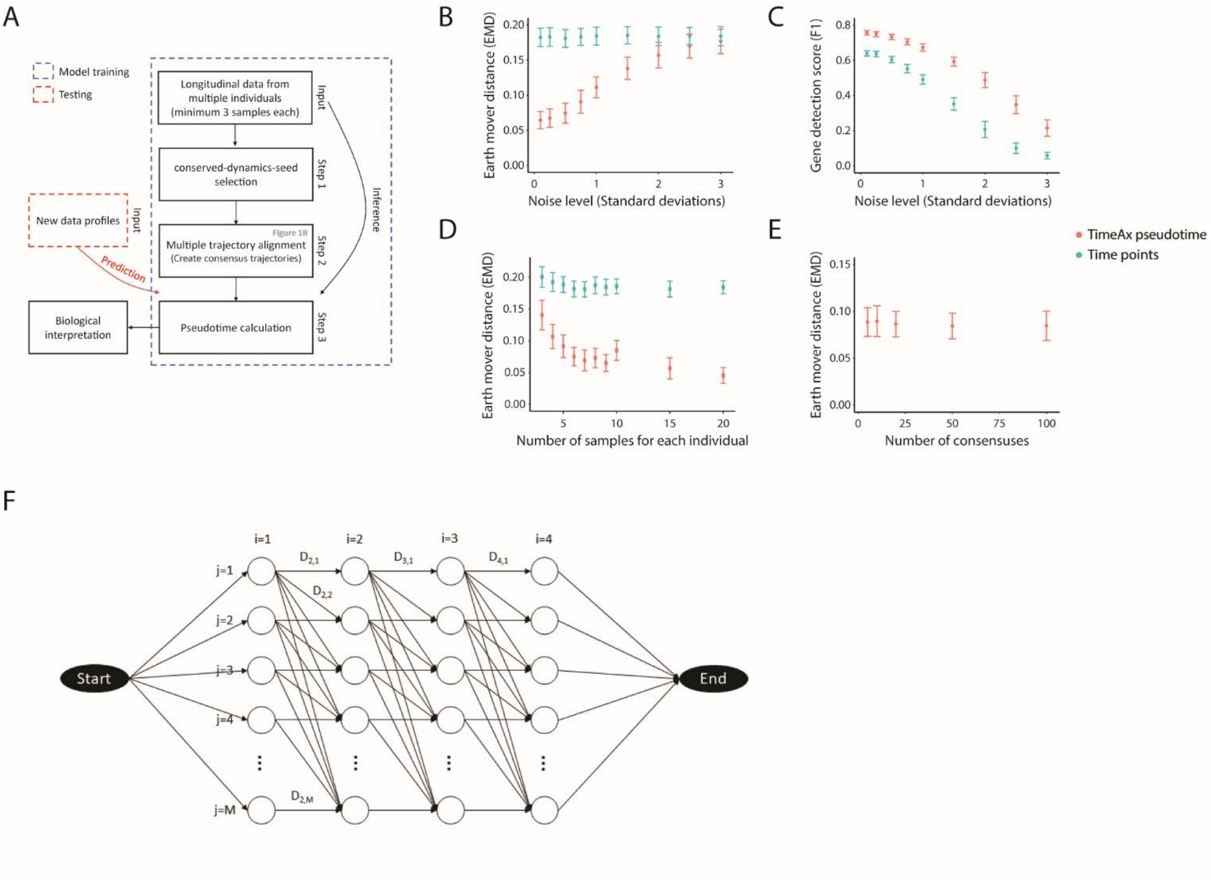


**Figure S1: A.** A workflow diagram of TimeAx. **B-E.** Simulated data analysis. Superior results for TimeAx predicted disease pseudotime (red), compared to simulated time points (blue), in capturing the the values of the simulated disease pseudotime **(B,D-E)** and for detecting dynamics-related features **(C)**. The measured accuracy is presented across different levels of noise **(B-C)**, number of samples for each individual **(D)** and number of consensus trajectories **(E)** (*x*-axis). Disease pseudotime In B,D-E, prediction accuracy was calculated using the EMD distance to the simulated disease pseudotime, while in C, gene detection accuracy was inferred using the F1 score (See **Methods**). For all panels, error bars represent mean ± std. **F.** An example graph for calculating the disease robustness pseudotime for an individual with 4 time points and a consensus trajectory with M disease pseudotime positions (See **Methods**).

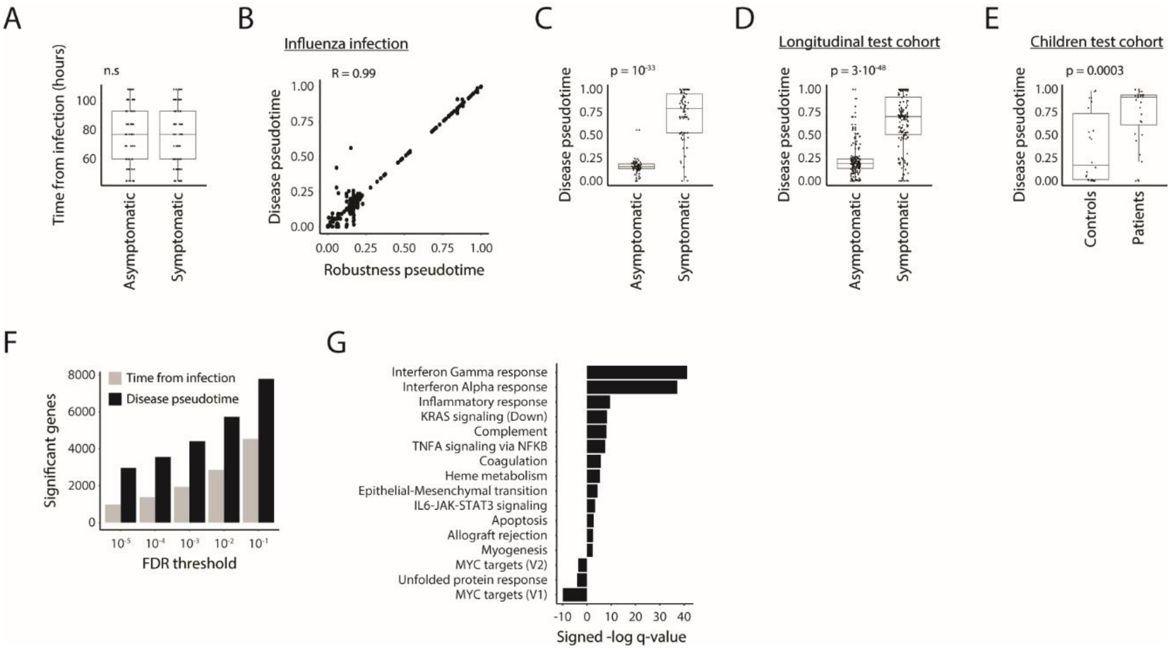


**Figure S2: A.** No significant difference in chronological time after infection (*y*-axis) between symptomatic and asymptomatic patients, in late time points after infection (>45 hours). **B.** TimeAx displays a high robustness score for the influenza infection model (r=0.99), calculated as the association between disease pseudotime (*y*-axis) and disease robustness pseudotime (*x*-axis, See **Methods**). **C-D.** Significantly higher disease pseudotime positions (*y*-axis) for symptomatic patients compared to asymptomatic patients, in late time points after infection (>45 hours), using the longitudinal train cohort (**C**) and the longitudinal test cohort (**D**). **E.** Significantly higher disease pseudotime positions (*y*-axis) for patients compared to healthy controls, using the children test cohort. **F.** Number of progression-related genes (*y*-axis), across different *q*-value thresholds (*x*-axis), is higher using pseudotime (black), compared to sampling time (gray), for the longitudinal train cohort (See **Methods**). **G.** Enrichment scores of biological pathways from MSigDB Hallmarks, based on associations of genes with the disease pseudotime in the longitudinal train cohort (See **Methods**).

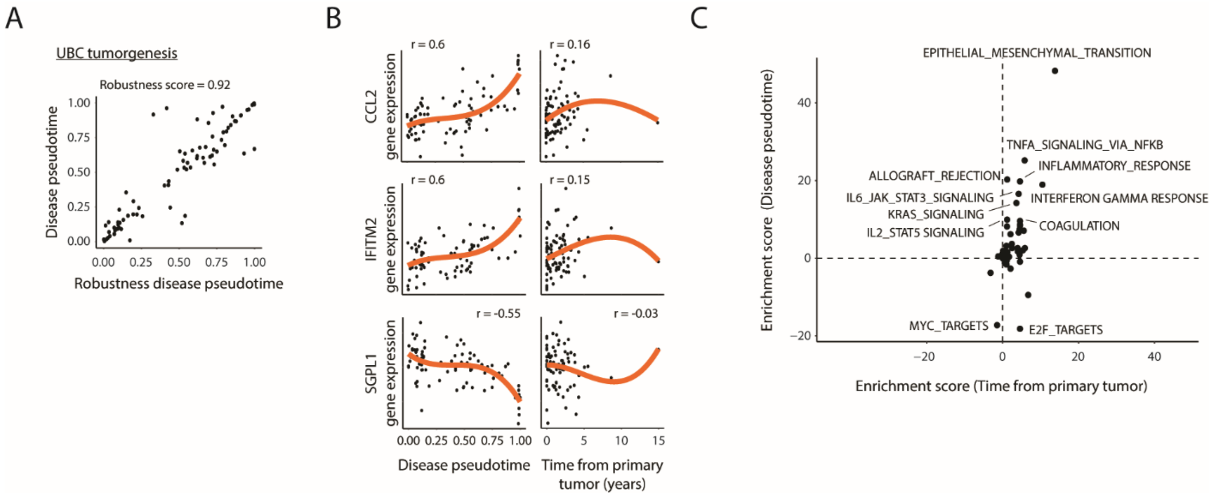


**Figure S3: A.** TimeAx displays a high robustness score for the UBC progression model (r=0.92), calculated as the association between disease pseudotime (*y*-axis) and disease robustness pseudotime (*x*-axis, See **Methods**). **B.** CCL2, IFITM2 and SGPL1 expression levels (*y*-axis) along either disease pseudotime (left; strong associations) or tumor recurrence times (right; weak associations) (*x*-axis). Regression trend line is displayed in orange. **C.** Enrichment scores of biological pathways from MSigDB Hallmarks (black dots), displaying improved scores while calculated using pseudotime (*y*-axis) compared to tumor recurrence times (*x*-axis).

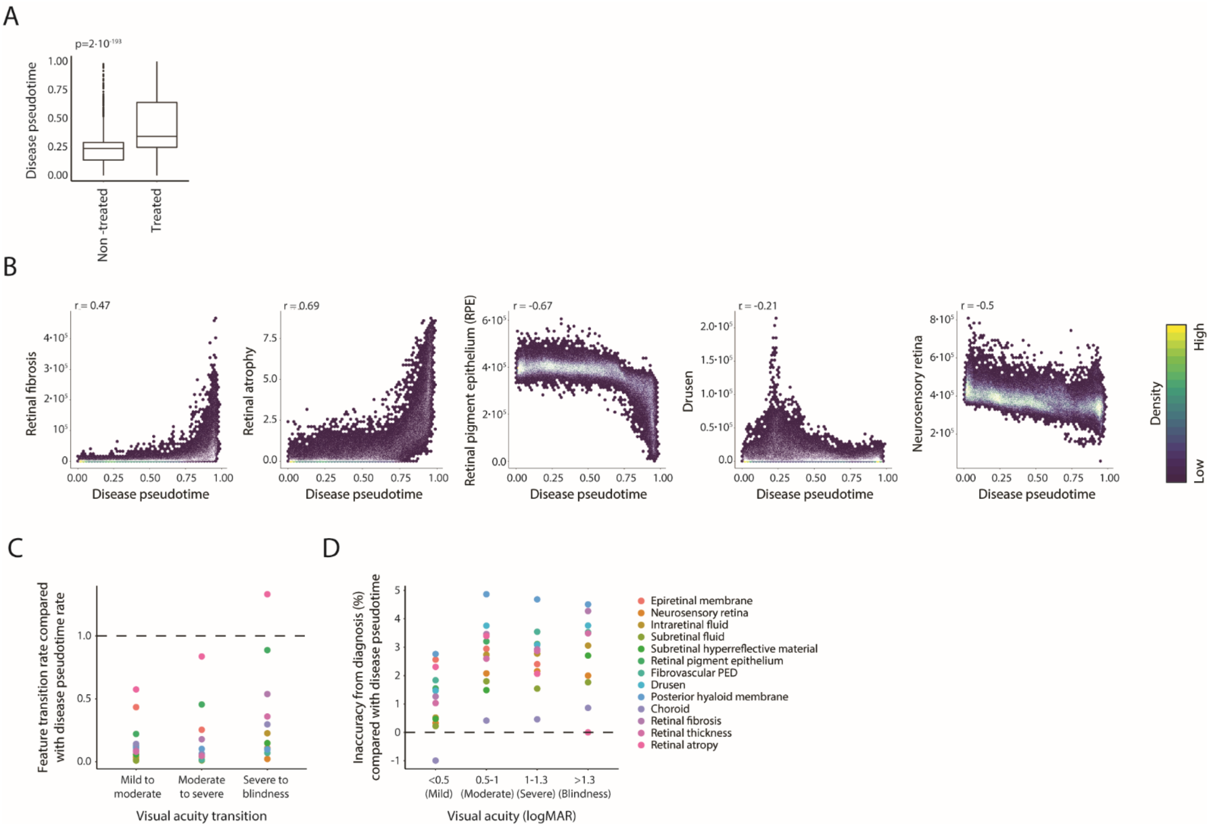


**Figure S4: A.** Disease pseudotime (*y*-axis) distribution in anti-VEGF treated and non-treated patients (*x*-axis) from the AMD test cohort. Boxes represent the 25th, 50th, and 75th percentiles; whiskers show maxima and minima. **B.** Associations of different segmented structural features of the retina (*y*-axis) with the disease pseudotime (*x*-axis), across all data points from the AMD test cohort. **C.** Ratios between segmented features' (color-coded) and disease pseudotime's average transition rates (*y*-axis), across each pair of consecutive visual acuity clinical states (*x*-axis) (See **Methods**). **D.** Percentage difference between segmented features' (color-coded) and disease pseudotime's error rates (*y*-axis), within each visual acuity clinical state (*x*-axis) (See **Methods**).

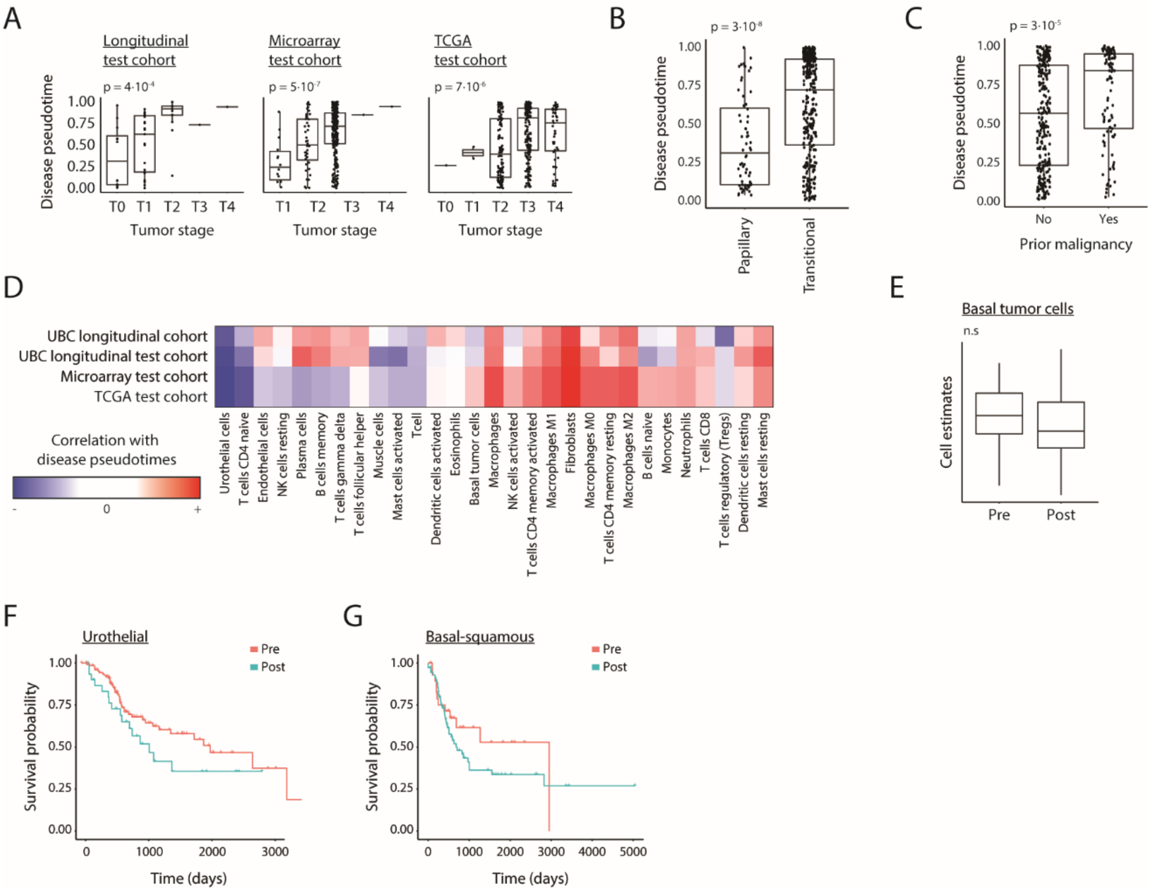


**Figure S5: A.** Shown are disease pseudotime (*y*-axis) relations with tumor stage (*x*-axis) for the longitudinal (left), microarray (middle) and TCGA (right) test cohorts. **B.** Disease pseudotime (*y*-axis) is significantly higher in transitional compared to papillary tumors. **C.** Disease pseudotime (*y*-axis) is significantly higher in patients with prior malignancies. **D.** Pearson correlations of deconvolved cell compositions with disease pseudotime, across all samples in both the UBC longitudinal train cohort as well as the longitudinal test, microarray and TCGA test cohorts (See **Methods**). Negative to positive correlations are colored in a blue to red color scale. **E.** Basal tumor cells deconvolved composition (*y*-axis) in pre- and post-inflection point samples (*x*-axis), presenting no significant difference between the two. **F-G**. Survival plots for either Urothelial-like (**F**) or Basal-squamous (**G**) patients within the TCGA validation cohort, comparing patients' tumors with disease pseudotime positions lower and higher the inflection point (pre vs. post, respectively; color coded). In **A**-**C** and **E**, boxes represent the 25th, 50th, and 75th percentiles; whiskers show maxima and minima.

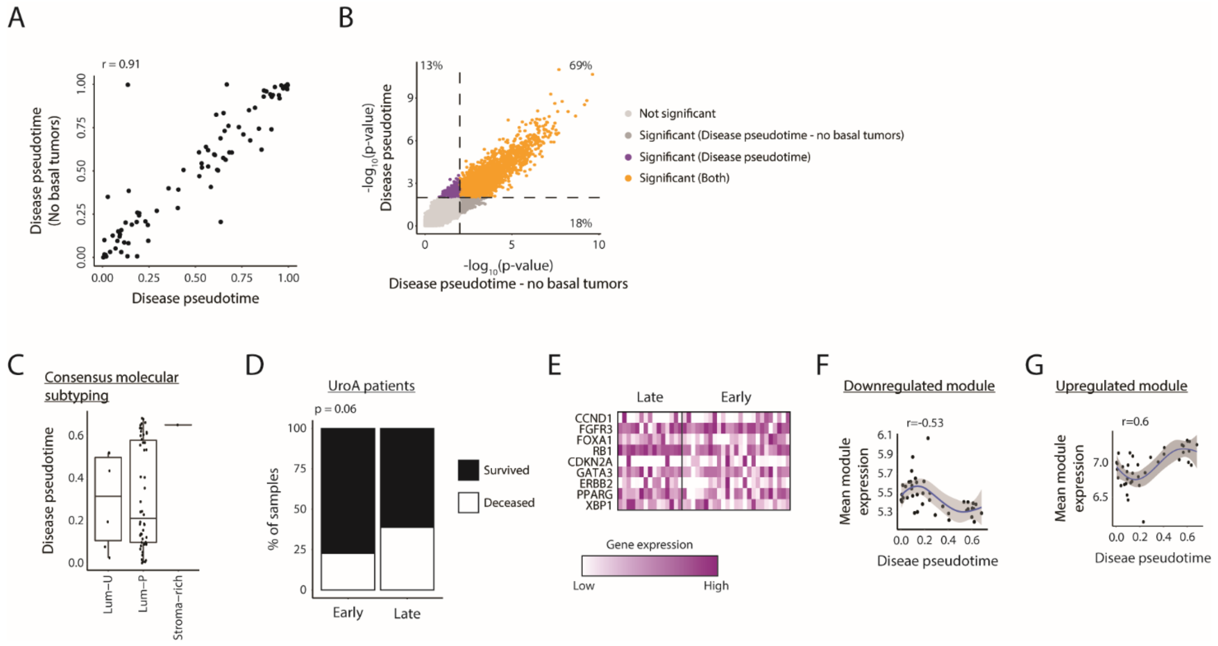


**Figure S6: A.** Strong association between disease pseudotime inferred by model using all patients (*x*-axis) and all patients with non-basal tumors (*y*-axis) from the UBC longitudinal cohort. **B.** Gene associations (-log10 transformed, FDR-corrected, q-values) with disease pseudotime inferred by model using all patients (*x*-axis) or all patients with non-basal tumors (*y*-axis), using a *p*-value threshold of $10^{-2}$ (Dashed lines) (See **Methods**). Genes are colored based on their association with the two time axes. **C.** Disease pseudotime distribution across tumor molecular classifications in patients pre-stromal pro-invasion point within the UBC longitudinal cohort based on the 'consensus' molecular subtyping framework. **D.** Survival (black) versus deceased (white) rates of UroA patients within the TCGA test cohort at the early and late patient groups (*x*-axis). p-value was calculated using Fisher's exact test. **E.** Heatmap of the expression levels of UroA marker genes (rows), in early and late UroA tumors (columns) within the UBC longitudinal cohort. **F-G.** Average expression levels of genes within the downregulated **(F)** and upregulated **(G)** modules (*y*-axis) along the disease pseudotime (*x*-axis) in patients pre-stromal pro-invasion point.

**Supp. Table legends**

**Table S1:** Statistics of the Cox proportional-hazards model classifying patients survival rates for either pre- or post- stromal pro-invasion point tumors, after accounting for other covariates with known association with survival, including age, sex and the clinical stage of the disease.

**Table S2:** Differential expression of genes (column 1) between UroA early tumors (mean expression in column 2) and UroA late tumors (mean expression in column 3). Significance value, based on student's t-test is presented in column 4..

**Table S3:** Highly enriched pseudogene and microRNAs groups within the downregulated module (column 1), their number of appearances in the module (column 2) and their equivalent enrichment score (*q*-value) within the module, based on a hypergeometric test (column 3).

**Table S4:** Highly enriched gene sets related to membrane channel proteins and GPCRs within the downregulated module (column 1), their number of appearances in the module (column 2) and their equivalent enrichment score (*q*-value) within the module, based on a hypergeometric test (column 3).

**Table S5:** Highly enriched gene sets within the upregulated module (column 1), their number of appearances in the module (column 2) and their equivalent enrichment score (*q*-value) within the module, based on a hypergeometric test (column 3).
